# The role of the neutrophil receptor Mrgpra2 in the formation of itch in atopic dermatitis

**DOI:** 10.64898/2026.06.12.731726

**Authors:** Taylor Follansbee, Henry Le Chang, Yanni Larsen, Ryo Kawamoto, Shekhar Pandey, Xinzhong Dong

**Affiliations:** The Solomon H. Snyder Department of Neuroscience, Johns Hopkins University School of Medicine, Baltimore, MD 21205, USA; Department of Neurosurgery, Johns Hopkins University School of Medicine, Baltimore, MD 21205, USA; Department of Dermatology, Johns Hopkins University School of Medicine, Baltimore, MD 21205, USA; Howard Hughes Medical Institute, Chevy Chase, MD 20815, USA

**Author notes:** Authors contributed equally (Taylor Follansbee, Henry Le Chang).

**Keywords:** Mrgpra2, MC903, Chronic itch, Tslp, Inflammation

## Abstract

Atopic dermatitis (AD), or eczema, is estimated to affect more than 30 million people in the United States, with over 6 million have moderate or severe disease. Chronic pruritus is a common symptom and is one of most difficult to manage. Even though itch is a major component in the pathology of AD, the biological underpinnings are not fully understood. In the present manuscript we identify a role for the neutrophil receptor, Mrgpra2, in the development of AD hyperplasia and chronic itch in the mouse. The role of Mrgprs in the context of itch have provided a huge step in our understanding of pruritus and have led to the development of novel therapeutics. Here we provide new evidence for the involvement of Mrgpra2 in the development of AD. Here we show that genetic deletion of Mrgpra2 significantly reduces scratching behaviors, transepidermal water loss, epidermal thickening, and Tslp expression in AD.

## INTRODUCTION

Atopic dermatitis (AD), or eczema, affects more than 30 million people in the United States, with over 6 million having moderate or severe disease. Chronic pruritus is a common symptom and is one of most difficult to manage since it disrupts sleep, negatively impacts mental health, and drives scratching behaviors which can further damage the skin barrier (Mollanazar et al., 2016). Even though itch is a major component in the pathology of AD, the biological underpinnings are not fully understood. The current study focuses on understanding the role of neutrophils, and more specifically the role of the mas-related G protein coupled receptor A2 (MrgprA2), in the development of atopic dermatitis.

Atopic dermatitis is classically considered a T-cell mediated disease, however recent evidence suggests an emerging role of neutrophils in the development of AD (Chiang et al.,2024; Choy et al., 2012). Transcriptomic analysis of human AD lesions identified neutrophil-associated inflammatory transcripts, implying that neutrophil mediated inflammation may contribute to disease (Choy et al., 2012). Mechanistically, it was found that neutrophils infiltrate the skin in the MC903 murine AD model and sensitize pruriceptors via release of Cxcl10, resulting in increased scratching behavior (Li et al., 2006; Walsh et al., 2019). Inhibition of neutrophil infiltration significantly reduced scratching behaviors and reduced pruriceptor sensitivity (Pavlenko et al., 2023), further supporting a role for neutrophils in the development of AD.

Mrgprs are a family of receptors well known for their role in itch signaling and immune cell function (Dong et al., 2022; Gour et al., 2024; Liu et al., 2009). Neutrophils selectively express the duplicate genes for Mrgpra2a and Mrgpra2b, collectively referred to as Mrgpra2, and amongst granulocytes, expression of Mrgpra2 is strictly limited to neutrophils. Activation of Mrgpra2 by anti-microbial peptide, defensin 14, induced neutrophil recruitment and abscess formation during bacterial skin infection (Dong et al., 2022). Double knockout of Mrgpra2a/b (A2KO) resulted in a significant reduction in the number of neutrophils to the site of infection. Despite the increasing recognition of neutrophils in atopic dermatitis, whether Mrgpra2 contributes to the development of AD has not been explored. In this study we tested whether Mrgpra2 contributed to the development of AD in the mouse model.

## RESULTS

Mice received daily application of MC903 for seven consecutive days. Consistent with previous reports, we observed a significant time-dependent increase in scratching behaviors (Fig. 1).

**Figure 1.**
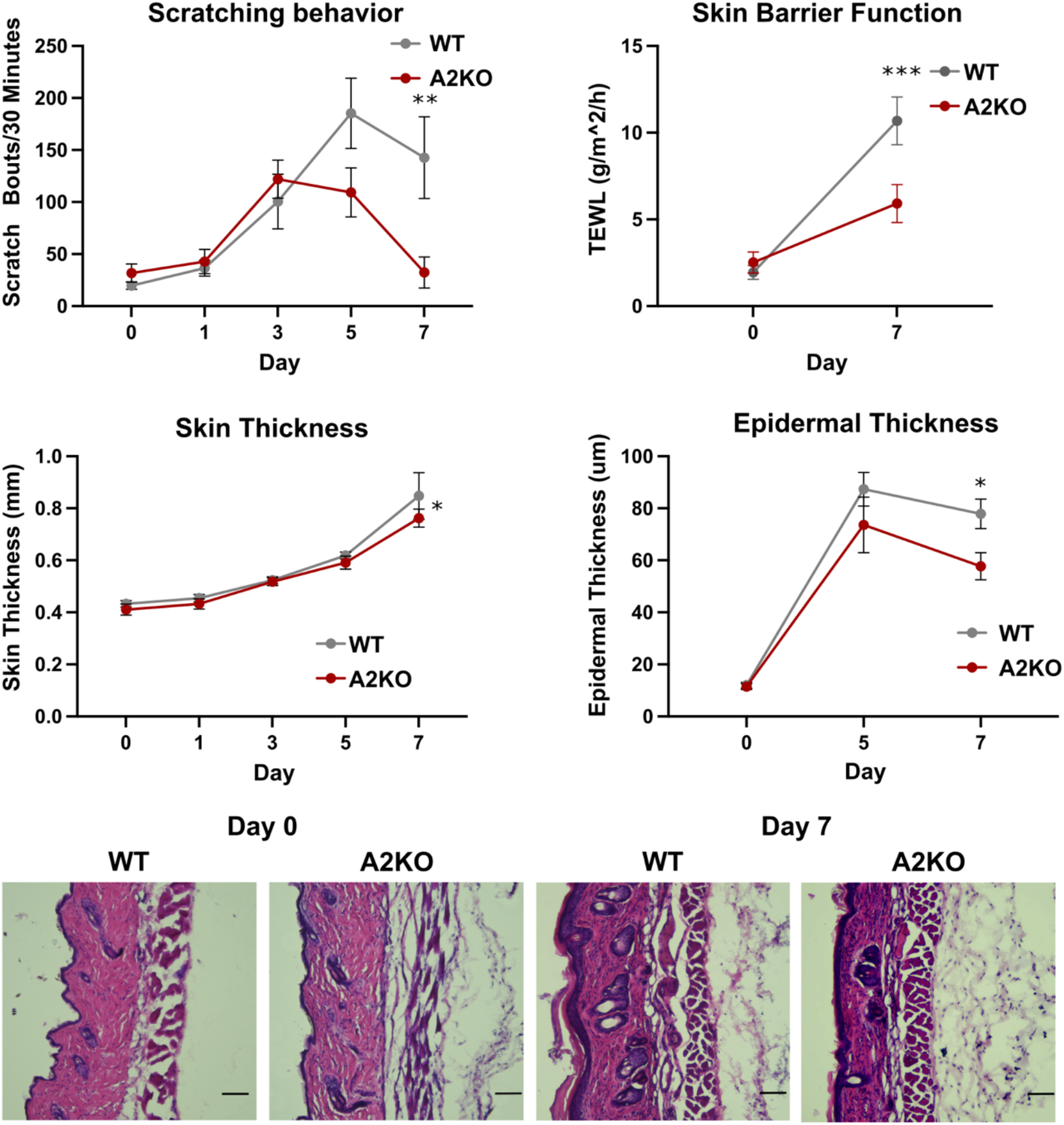
Skin pathology and behavior in MC903 model of atopic dermatitis. Daily treatment with MC903 resulted in increased scratching behaviors (top left; WT, n = 5; A2KO, n = 7) and increased trans-epidermal water loss (TEWL; top right; WT, n = 13; A2KO, n = 13), which was significantly reduced in A2KO mice. MC903 treatment induced skin (middle left; WT, n = 4-10; A2KO, n = 4-8) and epidermal thickening (middle right; WT, n = 5-13; A2KO, n = 5-8), which was reduced in A2KO mice. Representative images of epidermal skin thickening from WT and A2KO mice on day 0 and day 7. Two-way ANOVA with Sidaks Multiple comparisons, *p<0.05, **p<0.01, ***p<0.001. Scale bars = 100 um.

Compared to WT mice, we observed a significant decrease in the scratching behaviors of A2KO mice on day 7 and a non-significant decrease on day 5 (Fig. 1). Similarly, Mrgpr-cluster knockout mice (Liu et al., 2009), in which a cluster of Mrgpr genes including Mrgpra2a/b are deleted, had significantly fewer scratch bouts on Day 7 compared to WT controls (Fig. S1). Treatment with ethanol vehicle did not induce time dependent changes in scratching behaviors for WT or A2KO mice (S1), however baseline scratching behaviors were higher in A2KO mice when compared to WT mice.

In addition to the behavioral changes, we observed significant effects of both day and genotype in the total skin thickness, a pathology of AD (Chiang et al., 2024), with the skin thickness progressively increasing over the course of treatment for WT and slightly less thickening in A2KO mice. While multiple comparisons testing did not identify any significant timepoint differences between genotypes, we did observe a significant decrease in epidermal thickness of A2KO mice on day 7 (Fig. 1). To further evaluate skin barrier dysfunction, we measured trans-epidermal water loss (TEWL) on days 0 and 7 of MC903 treatment and found a significant increase in TEWL following MC903 treatment, which was significantly reduced in A2KO mice (Fig. 1). Taken together, these results demonstrate that loss of Mrgpra2 reduces chronic itch related scratching behaviors and epidermal barrier dysregulation in the MC903 AD model.

Given the role of Mrgpra2 in neutrophil recruitment during bacterial skin infection, we sought to determine whether the reduced itch and epidermal pathology observed in A2KO mice was due to reduced neutrophil skin infiltration. Ly6G positive neutrophils were histologically counted on days 0, 5, and 7 of treatment, correlating to the days with the most robust changes. Infiltrating neutrophils in the skin were significantly increased following treatment with MC903 (Fig. 2), confirming substantial neutrophil recruitment during treatment, however no significant differences in neutrophil count were observed between WT and A2KO mice. To further investigate infiltrating neutrophils, we performed flow cytometry. Relative to CD45 immune cells, the proportion of infiltrating neutrophils was significantly increased during disease progression, but no differences between WT and A2KO mice were observed (Fig. 2).

**Figure 2.**
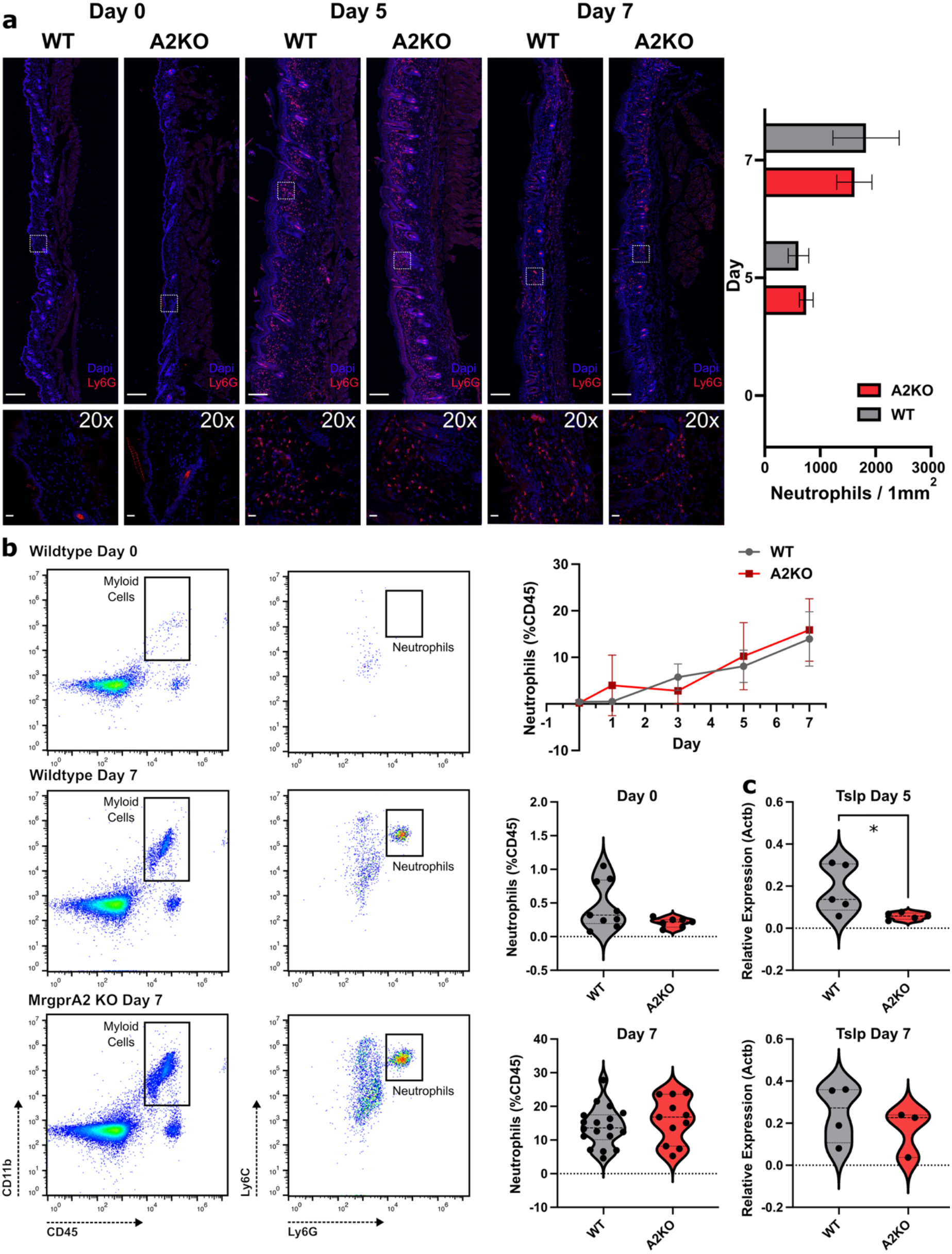
Skin immune cell recruitment in the MC903 model of atopic dermatitis. Infiltrating neutrophils in the skin were quantified histologically with Ly6G staining on days 0, 5, and 7. Neutrophil count was normalized to 1mm^2^ and WT and A2KO mice were directly compared (WT, n = 5-9; A2KO, n = 4-5, Two-way ANOVA with Sidaks Multiple comparisons). Flow cytometry was used to determine the proportion of infiltrating neutrophils (neutrophils/CD45+ cells) during MC903 treatment (WT, n = 3-18; A2KO, n = 3-11, Two-way ANOVA with Sidaks Multiple comparisons). Tslp RNA expression was measured from skin samples on day 5 and day 7 for WT and WTKO mice (WT, n = 4-5; A2KO, n = 3-5, Students t-test). *p<0.05, **p<0.01, ***p<0.001. Scale bars for whole skin images = 200 um, and 20X images = 20 um.

Additionally, neutrophil elastase in Ly6G positive skin cells was not significantly different between genotypes on days 5 and 7 (Fig. S2).

Finally, we investigated expressional changes of inflammation-related genes in the skin of MC903 treated WT and A2KO mice. We found that Tslp, a hallmark inflammatory cytokine associated with AD (Wilson et al., 2013), was significantly reduced in A2KO mice on day 5 of treatment, but not day 7 (Fig. 2). Several additional neutrophil and inflammatory related genes were examined. Relative expression of Il1b, Cxcl10, Cxcr2, and Cxcr4 were not significantly changed (Fig. S3). While none of these targets were independently significant, there was a consistent reduction across these gene targets particularly on day 5. Our results suggest that loss of Mrgpra2 may alter neutrophil inflammatory signaling early in the development of AD.

## DISCUSSION

Our findings support a role for Mrgpra2 signaling in the development of chronic pruritus and epidermal barrier dysfunction in the murine MC903 AD model. Surprisingly, despite the robust reduction in scratching behaviors and improvement in skin barrier function which we observed in A2KO mice, neutrophil recruitment was not changed. Our findings suggest that Mrgpra2 may regulate neutrophil activation or signaling rather than infiltration. The reduction in Tslp, as well as general decreased inflammatory cytokines, observed on day 5 but not day 7 supports neutrophils as an early disease mediator for AD progression. Future studies are needed to determine the mechanism underlying Mrgpra2 mediated inflammation in AD which may lead to novel therapeutic targets.

## ETHICS STATEMENT

This study was performed in accordance with our approved ACUC protocol at Johns Hopkins University.

## DATA AVAILABILIITY STATEMENT

Data will be made available upon request.

## CONFLICT OF INTEREST STATEMENT

We have no conflict of interests to report.

## ACKNOWLEDGEMENTS

This work was supported by the Howard Hughes Medical Institute and R37NS054791 (X. Dong) and the NIH T32NS070201 (T. Follansbee) and the Woodrow Wilson Undergraduate Research Fellowship (H. Le Chang).

## METHODS

### Mouse behavior

Experiments were performed using wildtype mice (c57BL/6J, Jackson Labs, Bar Harbor ME), Mrgpra2a/b double knockout mice (A2KO), and Mrgpr Cluster Knockout mice (MrgprA1–A4, A10, A12, A14, A16, A19, B4, B5, and C11). Mice of both sexes, aged 8-12 weeks old, were used. Mice were given free access to water and housed in standard animal housing facilities, unless being used immediately for behavior or for tissue extraction and were kept on a 12 hour light/dark cycle. Mice were habituated for at least 3 days to the behavior apparatus and shaved 3 days prior to the start of treatment. All procedures were approved by the Johns Hopkins ACUC (protocol?). Scratching behaviors were recorded on days 0, 1, 3, 5, 7 prior to application with 40 uL MC903 (20 uM in EtOH; calcipotriol, TOCIS, 2700) to the rostral back. Mice were video recorded from below and allowed to habituate for 15 minutes after being placed in the glass cylinders. Recorded videos were analyzed from 2-unbiased observers.

### Skin pathology

Skin barrier function was assessed by measuring transpidermal water loss (CyberDERM) on day 0 prior to the start of treatment and on day 7 of treatment. Total skin thickness was measured on days 0, 1, 3, 5, 7 using calipers at the center of treatment area.

### Immunofluorescence staining and microscopy

Skin sections were biopsied from euthanized mice using an 8mm skin punch and postfixed in 4% PFA for 1 day, followed by 2-3 days in 30% sucrose. Samples were then frozen in OCT for cryo-sectioning at 12 or 20 um thickness. Skin sections were mounted to slides and frozen at -20 C until use. Skin sections were stained with H&E (Hematoxylin and Eosin Stain Kit, Vector Laboratories) and the slides sealed with Cytoseal. For immunofluorescence staining, slides were washed with PBST (0.1%) for 5 minutes and blocked in 10% NGS for 1 hour. Primary antibody (Ly6G, Bio X Cell) was applied at 1:1000 overnight at 4C and secondary (568 or 488 – Goat anti-rat, 1:500) was applied for 2 hours at room temperature. Elastase was stained using anti-Neutrophil Elastase (Thermo Fisher Scientific) at 1:500 dilution in 5% NGS, with a 568 – Goat anti rabbit secondary at 1:1000 dilution. Slides were mounted with Fluoromount-G with DAPI (Invitrogen). Images were taken using a Zeiss confocal microscope and analyzed with FIJI.

### Flow cytometry

Skin punch biopsies (8 mm) were finely cut and digested in 5 mL of digestion buffer containing 100 mg/mL DNase I (Worthington Biochemical) and 1.67 Wunsch units/mL Liberase TL (Roche) in RPMI for 2 hours at 37 C. Single cell suspension was filtered through a 40 um strainer and washed in RMPI, followed by PBS. Live dead stain was applied for 15 minutes (Aqua Dead Stain, ThermoFisher). Cells were blocked for 5 minutes with CD16/CD32 Fc block (A170011E, Biolegend) and stained for 25 minutes with the following antibodies: CD45-APC/Cy7 (30-F11, Biolegend), CD11b-PE/Dazzle594 (M1/70, Biolegend), Ly6C-APC (HK1.4, Biolegend), Ly6G-BV421 (1A8, Biolegend). Neutrophils were identified as CD45+/Ly6G+/CDll1b+ and reported as a percentage of CD45+ cells. Sample data was acquired with a Cytoflex LC (Beckman Coulter) and analyzed with FlowJo (BP).

### Quantitative PCR

Snap frozen skin biopsies (8 mm) were homogenized in TriReagent and RNA was extracted using the Direct Zol RNA Miniprep Plus kit (Zymo). RNA was reversed transcribed into cDNA using the HiScript IV RT SuperMix for qPCR (Vazyme). qPCR was performed using the Taq Pro U^+^ Multiple Probe qPCR Mix (Vazyme) with FAM Taqman gene expression probes (Thermo Fisher). Cycling and data acquisition was performed with a Quantstudio 5 PCR system (applied biosystems).

### Statistics

All statistical analyses were performed using Prism 11 (Graphpad). Single comparisons were made using two-tailed students t-tests and multiple comparisons were made using two-way ANOVAs with posthoc testing.

## Notes

### Competing Interest Statement

The authors have declared no competing interest.

