## Supplemental Files for "The role of the neutrophil receptor Mrgpra2 in the formation of itch in atopic dermatitis"

### SUPPLEMENTARY FIGURES

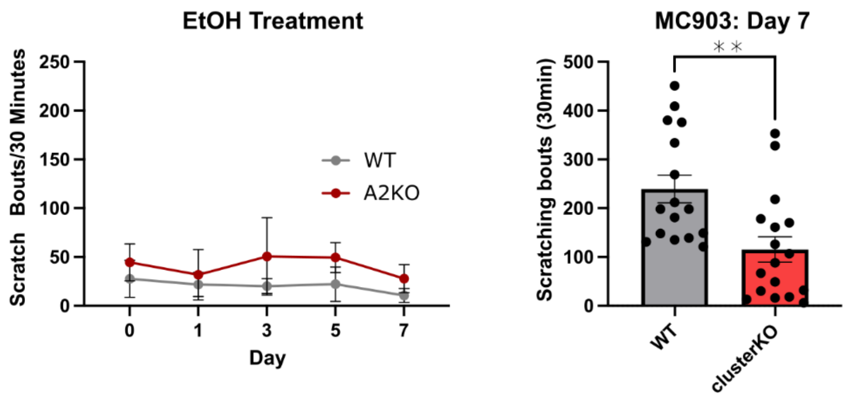


#### Supplementary Figure 1

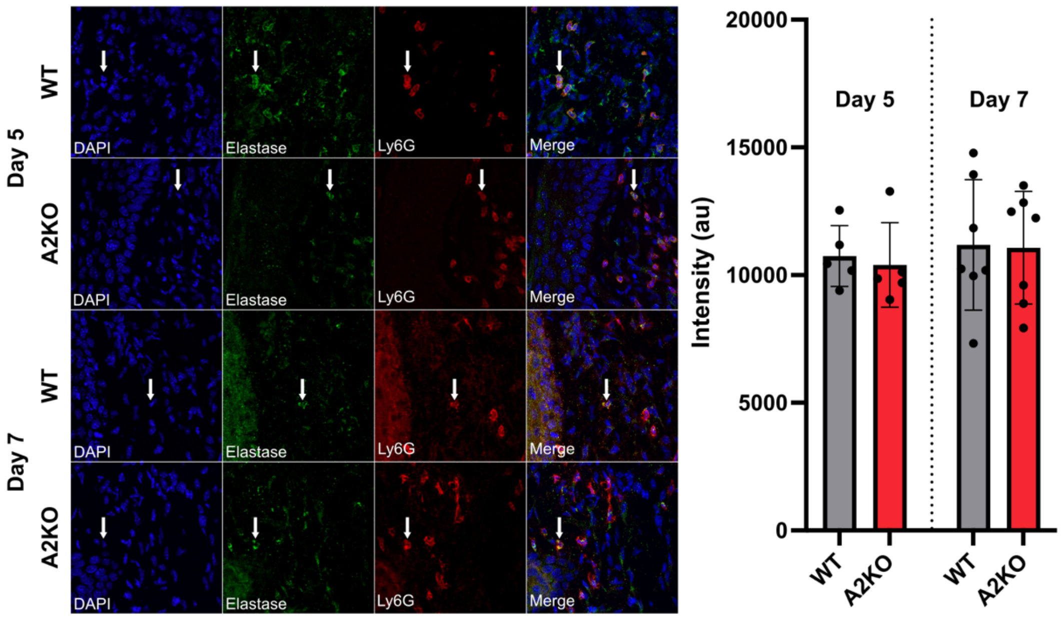


#### Supplementary Figure 2

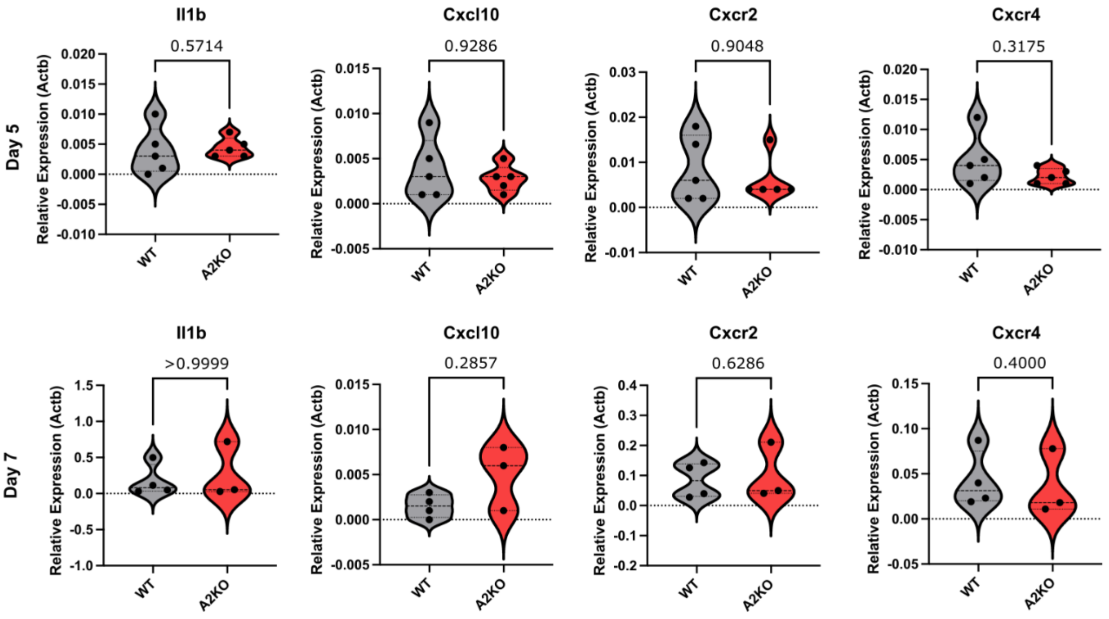


**Supplementary Figure 3**

**Supplementary Figure S1. Scratching behavior in WT and knockout mice.**

Scratching bouts in WT and A2KO mice during daily ethanol vehicle treatment, left. Two-way ANOVA with Sidaks Multiple comparisons (WT, n = 8; A2KO, n = 5) Scratching bouts in WT and Mrgpr-cluster knockout mice on day 7 of MC903 treatment, right. Students T test (WT, n = 16; A2KO, n = 17). *p<0.05, **p<0.01, ***p<0.001.

**Supplementary Figure S2. Neutrophil elastase staining in MC903-treated skin.**

Representative images of WT and A2KO skin sections stained for elastase (green), Ly6G (red), and DAPI (blue) on days 5 and 7 after MC903 treatment. Arrows indicate representative Ly6G$^{+}$ elastase$^{+}$ cells. Right, quantification of elastase fluorescence intensity. Two-way ANOVA with Fishers LSD (WT, n = 5; A2KO, n = 7).

**Supplementary Figure S3. Inflammatory gene expression in MC903-treated WT and A2KO skin.**

Relative expression of *Il1b*, *Cxcl10*, *Cxcr2*, and *Cxcr4* in WT and A2KO skin after 5 or 7 days of MC903 treatment. Expression was normalized to *Actb*. Each point represents one mouse. No significant differences were observed between genotypes. (WT, n = 4-5; A2KO, n = 3-5).
